# Multi-locus CRISPRi targeting with a single truncated guide RNA

**DOI:** 10.1101/2023.10.20.563306

**Authors:** Molly M Moore, Siddarth Wekhande, Robbyn Issner, Alejandro Collins, Yanjing Liu, Nauman Javed, Jason D Buenrostro, Charles B Epstein, Eugenio Mattei, John G Doench, Bradley E Bernstein, Noam Shoresh, Fadi J Najm

**Author notes:** These authors contributed equally to this work (Molly M Moore, Siddarth Wekhande).

## Abstract

A critical goal in functional genomics is evaluating which non-coding elements contribute to gene expression, cellular function, and disease. Functional characterization remains a challenge due to the abundance and complexity of candidate elements. Here, we develop a CRISPRi-based approach for multi-locus screening of putative transcription factor binding sites with a single truncated guide. A truncated guide with hundreds of sequence match sites can reliably disrupt enhancer activity, which expands the targeting scope of CRISPRi while maintaining repressive efficacy. We screen over 13,000 possible CTCF binding sites with 24 guides at 10 nucleotides in spacer length. These truncated guides direct CRISPRi-mediated deposition of repressive H3K9me3 marks and disrupt transcription factor binding at most sequence match target sites. This approach is valuable for elucidating functional transcription factor binding motifs or other repeated genomic sequences and is easily implementable with existing tools.

## Main

Over 1 million human cis-regulatory elements (CREs) have been cataloged across various cell and tissue types ^1–4^. CREs include the promoters, enhancers, insulators, and silencers that direct gene expression, sometimes in dynamic interplay or synergy. CRE function is further influenced by cell state and multiple transcription factor (TF) binding sites. TFs recruit proteins and complexes to orchestrate gene expression. TFs bind with various strengths, often dictated by cell state and genomic contexts such as motif combinations and orientations ^5–7^. However, the determinants for TF binding to one motif over another and the effect of that binding are not well understood. Connecting CREs and TF binding with functional outputs is important for interpreting disease associated genetic variation^3,8,9^ and may help nominate regions for clinical interventions. Together, TFs and CREs direct the intricate regulatory networks that govern cell function and disease.

CRISPR interference (CRISPRi) consists of a catalytically dead Cas9 (dCas9) that can be fused to a zinc-finger repressive protein (KRAB) for transcriptional silencing. Several studies have relied on CRISPRi-directed targeting of CREs followed by RNA measurement or flow cytometry to detect gene expression changes ^10–19^. However, efforts to characterize CREs at scale have been complicated by the large number of putative elements and mild effect sizes. High multiplicity of infection (MOI) delivery of guides paired with single-cell RNA-seq provided a multiplexed testing approach ^16^, though at the cost of many viral integration events. As such, while CRISPRi-based approaches can effectively assess significant CREs, there is a critical need for improving their scalability.

The Cas9 nuclease is guided by a spacer sequence that determines targeting specificity. Typically, spacers are 20 nucleotides (nt) in length and target a single genomic site based on sequence complementarity. Early studies posited that spacers with minor 5’ truncations or mismatches retain Cas9-mediated, on-target cleavage ^20–22^. The 3’ end of the spacer sequence, also termed the seed sequence, is necessary though not alone sufficient for on-target cleavage. Activity was observed with truncated spacers of 17nt while 15nt or shorter spacers failed to demonstrate cleavage activity ^20,23–25^. However, an important distinction exists in the requirements for Cas9 binding and cleavage that is illuminated with dCas9 protein. Indeed, spacers as short as 10nt sufficed for dCas9-VPR (CRISPRa) activity at a single target site ^26^. We postulated that KRAB-dCas9 (CRISPRi) would perform similarly and, further, target multiple intended sites simultaneously.

Here, we explored the ability of truncated guides to direct CRISPRi to multiple sites simultaneously in the genome for multi-locus repression. Truncated guides resulted in reliable on-target efficacy down to spacer lengths of 9nt. TF motifs, which are often less than 14nt, presented ideal genomic loci for multiplexed repression. We target TF motifs in a CRE of the *EPB41* gene and observe comparable on-target efficiencies with full-length and truncated guides. We screened a truncated guide library targeting thousands of CTCF motif sites and discovered significant CTCF disruption. This approach offers a new opportunity to simultaneously perturb CREs at scale and effectively prioritize genomic loci for further study.

## Results

### Truncated guides direct CRISPRi to a sequence match site

We first set out to characterize the minimum guide length required for CRISPRi-mediated repression. CD81, a stably expressed, non-essential cell-surface protein, served as a reporter of on-target efficiency by flow cytometry (**Fig. 1a**). We selected a high performing 20nt *S. pyogenes* spacer (sgCD81i-1) ^27^, directed to the *CD81* transcriptional start site (TSS) and tested successive truncations. By convention, guides are cloned with a guanine in the 5’ position to improve Pol III transcription levels ^28^, sometimes resulting in the guanine complementing the target sequence (see **Methods** and **Supplementary Fig. 1a**). Therefore, here we use brackets to denote the length of guide sequence that complements a single target site. For example, sgCD81i-1 g[12nt] consists of a 5’ mismatched guanine and 12 complementary bases to the *CD81* TSS. Successive 5’ truncations of sgCD81i-1 resulted in repression with each guide down to a 9nt target match in Jurkat (T lymphocyte) cells (**Fig. 1b**), with sgCD81i-1 g[9nt] active and sgCD81i-1 g[8nt] exhibiting a complete loss of on-target activity. We next tested additional CD81 TSS 20nt guides with less effective on-target efficiency (sgCD81i-2 and sgCD81i-3). These truncated guides resulted in similar and sometimes better *CD81* repression relative to the respective 20nt guide (**Supplementary Fig. 1b**). We expected that CRISPR knockout would be ineffective with sizeable truncations based on prior studies ^23–25^ and designed 2 guides that target exon 1 of *CD81* (sgCD81-KO-1 and -2). CD81 knockout was effective at lengths down to a 17nt target match, consistent with prior findings (**Fig. 1c**). Indeed, guide length requirements for Cas9 cleavage and CRISPRi diverge at <17nt guide lengths, highlighting opportunities for CRISPRi targeting with truncated guides that are not possible with Cas9 cleavage.

**Fig 1.**
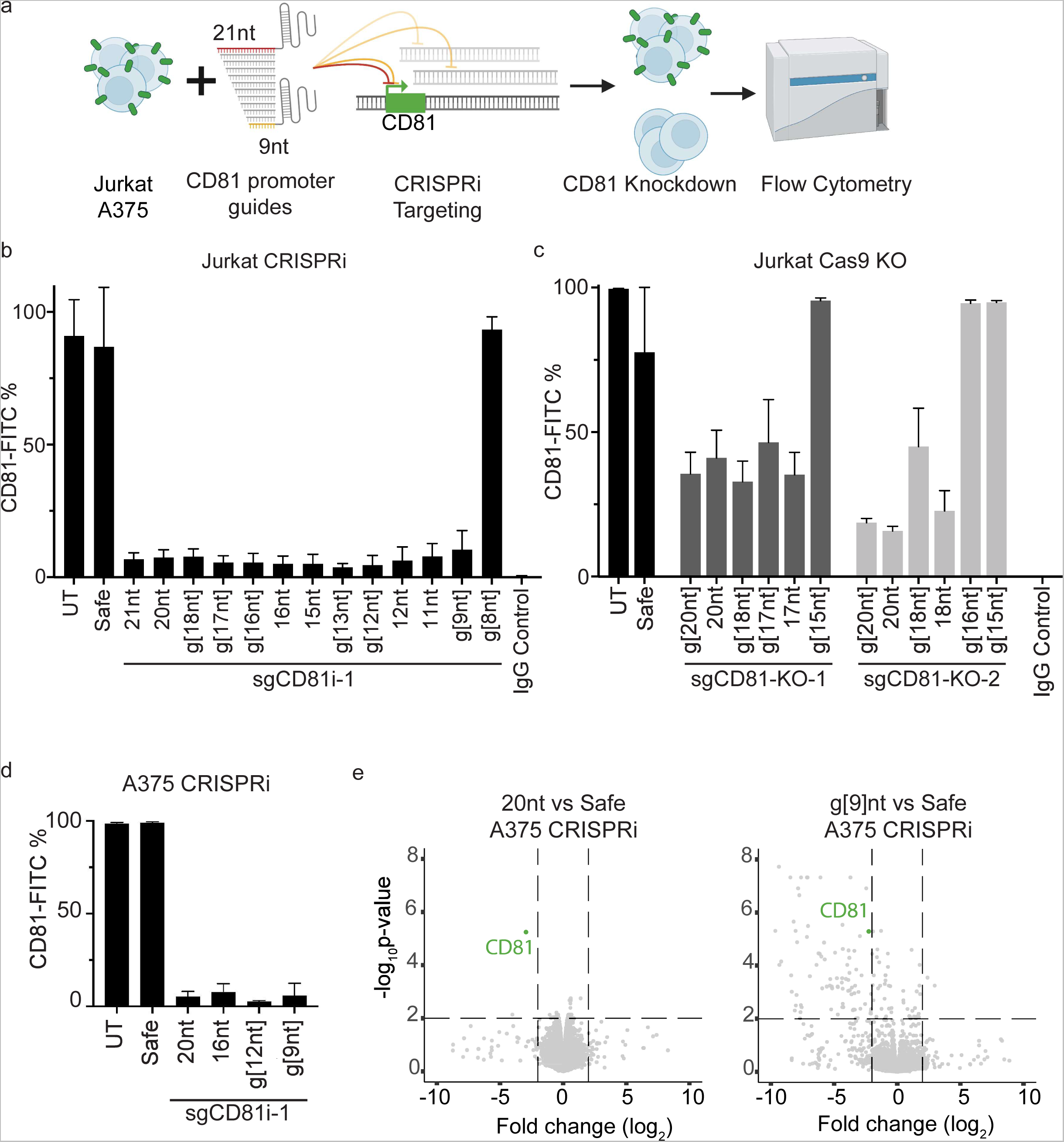
Truncated guides enable on-target CRISPRi-mediated repression. **a** Schematic of truncated guide experiments. **b** CRISPRi in Jurkat cells with truncated CD81 promoter targeting guide and analyzed for CD81 cell surface expression. **c** Cas9 cleavage in Jurkat cells treated with two guides targeting CD81 and truncated versions. **d** CRISPRi in A375 cells with truncated CD81 promoter targeting guide and analyzed for CD81 cell surface expression. **e** Gene expression in A375 cells in 20nt and g[9nt] sgCD81i-1 CRISPRi populations. p-value cutoff at >2 and log fold-change >2 and <-2. For **b-d**, cells were stained with CD81-FITC antibody analyzed by flow cytometry 7 days post lentiviral transduction for CD81 targeting guide, safe harbor control (Safe), or untransduced (UT). Flow gating strategy found in **Supplementary Fig. 1e**. Data are mean +/- SD from biological triplicate.

Next, we investigated the specificity of truncated guide repression. Unpaired bases at the 5’ end of 20nt guides can impact their activity. We lengthened the 5’ end of sgCD81i-1 10nt with 1-3 additional bases. Either 1 or 2 unpaired bases on the 5’ end resulted in effective repression, while 3 unpaired bases (gcc) completely abrogated repression (**Supplementary Fig. 1c, d**). We tested sgCD81i-1 full-length and truncated constructs in A375 (melanoma) cells and demonstrated similar *CD81* repression as observed in the Jurkat experiments (**Fig. 1d**), providing evidence that truncated guides are active in an additional cellular context. RNA sequencing in A375 showed similar levels of *CD81* repression at 20nt and g[9nt] lengths along with additional downregulated targets in the g[9nt] treatment (**Fig. 1e** and **Supplementary Table 1**). In sum, 5’ truncated guides can direct CRISPRi components to induce repression at target promoters comparable to full-length guides.

### Enhancer disruption with truncated guides

We next asked whether an active enhancer is targetable with truncated guides directed toward multiple TF motif sequences. We selected a 570bp locus with several putative TF binding sites, 2.8kb upstream of the *EPB41* gene (**Fig. 2a**, chr1:28,883,749-28,884,318). This locus was previously identified as a possible regulator of *EPB41* in a K562 CRISPRi screen ^16^. We selected 4 TF motifs in this enhancer (PU.1, SP1, YY1 and NR2), each containing an ideally positioned NGG sequence for the Cas9 protospacer adjacent motif (PAM) (**Fig. 2b**). TF motifs were positioned at the 3’ end of the guide including the PAM and two truncated versions (g[13nt] or 14nt and 11nt). The full-length guides match only the *EPB41* enhancer locus while the 11nt guides matched hundreds of additional genomic sites (**Fig. 2c**). We transduced K562 cells and measured on-target efficiency for *EPB41* knockdown by real-time quantitative PCR of *EPB41* and compared to 3 guides identified from the prior screen ^16^ as well as a promoter targeting guide (**Fig. 2d**). *EPB41* expression was reduced to levels comparable to the respective 20nt guide in 3 out of 4 11nt guides (PU.1, YY1, NR2) (**Fig. 2d and Supplementary Fig 2a**). In aggregate, the full-length and truncated guides tested significantly decreased EPB41 expression as compared to safe harbor control (**Fig. 2e** and **Supplementary Fig. 2b**, one-way ANOVA P<0.0001). CRISPRi-directed truncated guides can effectively disrupt an enhancer.

**Fig 2.**
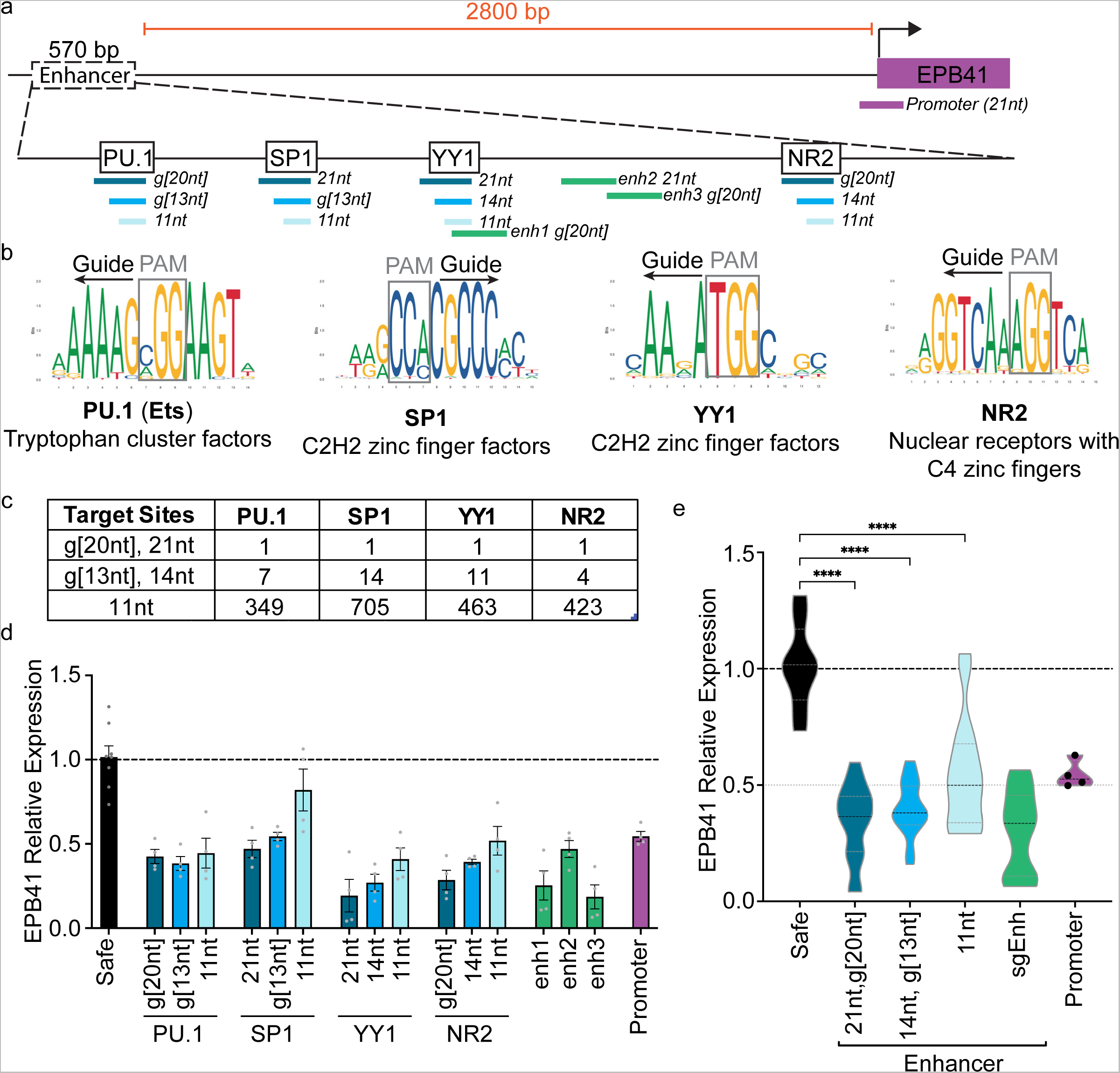
Enhancer targeting with truncated guides induces *EPB41* repression. **a** Schematic of *EPB41* genomic regulatory region and full-length/truncated guides targeting each of 4 TF motifs. **b** TF motifs (JASPAR) targeted by the guides with protospacer adjacent motif (PAM) orientation denoted. **c** Number of sequence match target sites in the genome (hg38) for the indicated motif directed guide. **d** Quantitative real time PCR analysis of *EPB41* expression in K562 cells with respective guide CRISPRi treatment conditions. Safe (Safe Harbor-targeting) guide = g[20nt], promoter guide = 21nt, enh1 = g[20nt], enh2 = 21nt, enh3 = g[20nt] (enhancer guides from Gasperini et al. 2019). Data are mean +/- SEM. **e** Violin plot summary of *EPB41* expression data in panel **d** organized by guide length or targeting region. TF-targeting guides (n=16; 4 guides per spacer length), sgEnh (n=12; 3 guides), and Promoter (n=4; 1 guide, all points shown) are the product of 2 biological replicates per guide run in duplicate. Safe is the sum of 3 biological replicates run 2-3 times each (n=8). Data are normalized to the mean CT value of Safe Harbor actin-b control probe. Results are analyzed by one-way ANOVA. ****P<0.0001 for all guide lengths compared with Safe Harbor guide.

### A CTCF-directed truncated guide library

To test the utility of truncated guides for multi-locus TF perturbation, we selected CCCTC-binding factor (CTCF) sites to screen. CTCF is a ubiquitously expressed TF whose role in genomic insulation is dependent on convergently oriented consensus sequences ^29^. Leveraging the 3’ NGG PAM sequence in the CTCF motif (**Fig. 3a)**, we designed a library of 24 10nt guides targeting a total of 13,352 sequence match CTCF binding sites (**Fig. 3a-c**). Based on CTCF ChIP-seq in Jurkat cells, approximately half of these sites are CTCF-bound (6,228) and represent 10.8% of all CTCF peaks (**Fig. 3c** and **Supplementary Fig. 3a**). This library allowed us to test CTCF binding sites, partitioned by guide, ranging from a minimum of 182 sites (sg1) to maximum of 1,123 sites (sg24). As a control we targeted the *CTCF* locus itself with full-length guides for gene repression (**Fig. 3d** and **Supplementary Fig. 3c)**. We packaged guides into a lentiviral library, transduced Jurkat cells near an MOI of 0.5, and collected cells over 21 days.

**Fig 3.**
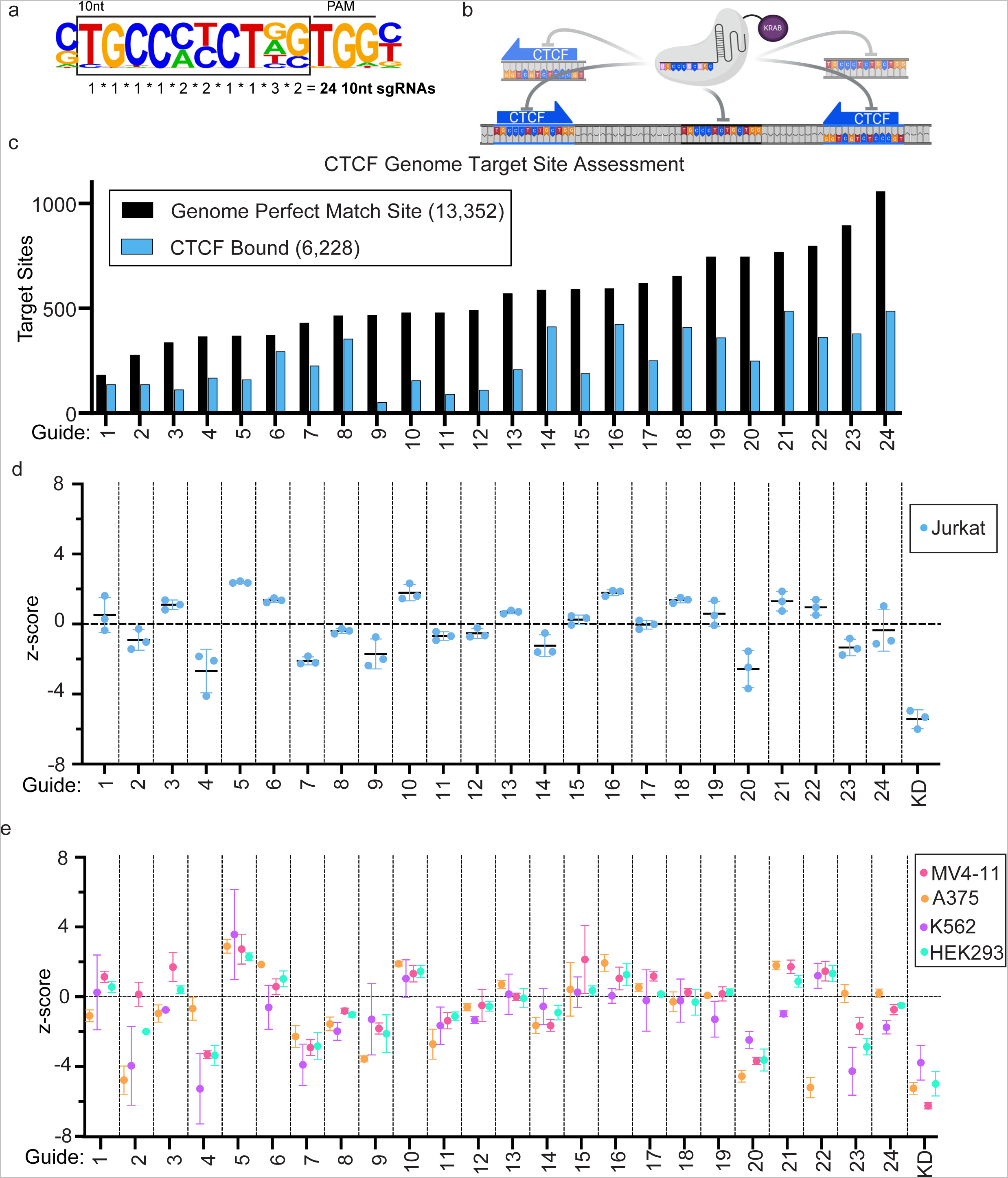
Simultaneous screening of CTCF binding sites with a truncated guide. **a** Guide selection strategy based on top Homer generated motif from CTCF ChIP in Jurkat cells. **b** CTCF targeting schematic of a single 10nt guide directing CRISPRi to many motifs simultaneously (CTCF-bound and unbound). **c** Target site distribution (black) and CTCF occupancy based on Jurkat ChIP (blue) for each of the 10nt guide sequences. **d, e** Guide enrichment and depletion in pooled screen for the indicated cell lines. Pooled screens were run in triplicate, while K562 was in duplicate. Data are mean +/- SD. CTCF promoter targeting full length guide knockdown (KD) used as a control.

We measured guide enrichment and depletion as a proxy for fitness. We quantified the scale of this effect with a z-score (see **Methods**) relative to 15 full-length safe harbor guides. Our results indicated that most truncated guides were not lethal, as we observed moderate shifts in guide representation relative to safe harbor guides (**Fig. 3d** and **Supplementary Tables 2 and 3**). A subset of guides resulted in enrichment, suggesting changes that may promote proliferation. We also identified guides sg4 and sg20 as broadly depleted, though not to the degree of *CTCF* knockdown. This initial screen provided evidence that certain truncated guides can induce fitness changes in Jurkat cells.

We next screened the CTCF library in additional cell lines to compare with Jurkat results. We processed A375 cells for CTCF ChIP-seq and found 6,140 library target sites bound, representing 14% of all CTCF peaks (**Supplementary Fig. 3a, b**). In comparison, 6,228 of CTCF library sites are bound in Jurkat. We further include K562 (T-lymphocytes), MV4-11 (AML) and HEK293 as additional models representing diverse cellular contexts for screening. Cells were transduced with the CTCF library and assessed for guide representation after 21 days. Fitness effects in these additional cell models largely recapitulated trends observed in Jurkat cells (**Fig. 3e** and **Supplementary Table 3**). This could be attributed to invariance of CTCF binding sites across tissues ^30,31^. However, we observed some instances of cell-specific fitness effects, particularly with sg2, sg22, and sg23. While sg2 and sg23 impacted more than one cell line, sg22 was strongly depleted in A375 only.

As an additional test of guide-sequence specificity, we screened 11nt guides by adding every base to each 10nt guide in the CTCF library, totaling 96 guides. The additional 5’ base had modest changes on fitness outcomes when compared to the respective 10nt guide outcome (pearson correlations ranged from 0.86 to 0.94 in Jurkat, 0.64 to 0.85 in K562, 0.79 to 0.94 in MV4-11, 0.86 to 0.96 in HEK293, and 0.73 to 0.87 in A375, **Supplementary Fig. 4**). This reinforced our prior finding that a single base mismatch was not detrimental to targeting, whereas ≥2 mismatches can disrupt activity (**Supplementary Fig. 1c-d**). Testing in 5 cell lines, we find that addition of a single 5’ base to a 10nt guide does not significantly alter guide effects.

### Simultaneous targeting of CTCF binding sites

We selected sg4 for further exploration due to its effects on fitness. Guide sg4 targets 357 sites in the genome with 10nt complementarity, termed “perfect match” sites. Perfect match sites were determined regardless of the 5’ guanine present on all guides. We transduced Jurkat cells with sg4 and CRISPRi for 6 or 7 days and performed CTCF and H3K9me3 ChIP-seq. The analysis revealed a significant drop in CTCF occupancy at perfect match sites (*t-test*, P<10^-5^) (**Fig. 4a, c** and **Supplementary Fig 5a, b**). We found it promising that CTCF, a strong binder to chromatin, was displaced at many sites simultaneously. Concurrent with CTCF loss was a significant increase in H3K9me3 signal at perfect match sites (*t-test*, P<10^-5^) (**Fig. 4b, d)**. H3K9me3 is a histone mark that indicates KRAB-dCas9 binding and recruitment of repressive proteins ^11^. Two example tracks of perfect match sites depict decreased CTCF binding and concurrent increased H3K9me3 signal (**Fig. 4e, f**). These initial findings presented compelling on-target CTCF disruption at multiple sequence match genomic loci with truncated guides.

**Fig 4.**
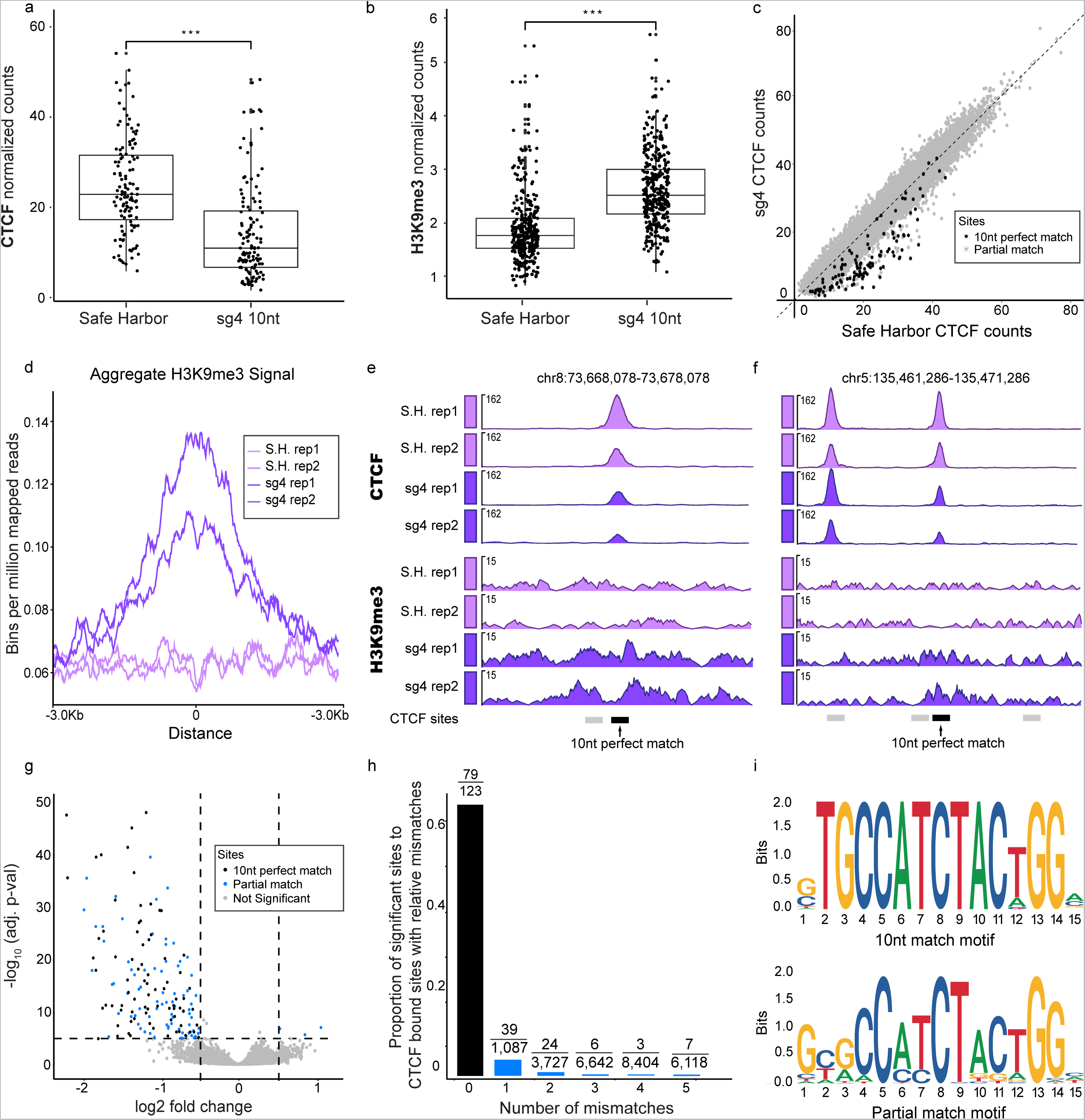
CRISPRi with a single truncated guide disrupts CTCF binding at multiple loci. **a** CTCF ChIP of the CTCF bound sites (357) targeted by sg4 with a perfect match. Significance with t-test ***P=5.6 x 10^-13^. **b** H3K9me3 ChIP of all sg4 perfect match sites. Significance with t-test ***P=1.3 x 10^-38^ . **c** Scatter plot CTCF binding at JASPAR motifs (880k) with 357 perfect match sites (black) and partial match sites (grey). **d** H3K9me3 density plot for Safe Harbor (S.H., light purple) and sg4 (dark purple) treated Jurkat cells. **e, f** Track views of 2 targeted regions depicting CTCF and H3K9me3 signal from 2 replicate experiments (rep). Bars on the bottom depict CTCF perfect match (black) and partial match sites (grey). **g** Volcano plot of CTCF bound sites (400k) with significant 10nt match CTCF sites (black), significant partial match sites (79), and not significant sites. Significance was determined at cutoffs (abs [LFC]>0.5 and −log_10_ P>5). **h** Histogram depicting significant CTCF disrupted sites as a fraction of CTCF bound sites with 0 mismatches up to 5 mismatches. Raw values depicted on each bar. **i** Logogram of significant perfect match sites (top, 79 sites) and significant partial match sites (bottom, 79 sites) from sg4 experiments.

We next evaluated multi-locus targeting specificity with a slightly longer spacer sequence. We designed a 13nt guide based on our prior findings that 3 mismatched bases disrupt targeting of *CD81* TSS (**Supplementary Fig. 1c-d**). We appended 3 bases (aag) to the 5’ end of sg4, termed aag[sg4], resulting in a guide with 77 expected perfect match sites. We transduced Jurkat cells with the guide and CRISPRi and processed cells for CTCF ChIP-seq. Examining the 26 CTCF-bound genomic regions targeted by both guides, we observed significantly lower CTCF signal in the aag[sg4] and sg4 samples compared to safe harbor guide (**Supplementary Fig. 6**). Sg4 and aag[sg4] were not significantly different from one another, however we observed a slightly lower mean CTCF signal in aag[sg4].

An additional truncated guide from the CTCF library was investigated further for its efficiency. We selected sg8 (10nt) which had 465 sequence match sites and little fitness impact on Jurkat cells. We transduced Jurkat cells with sg8 and CRISPRi and processed the cells for ChIP-seq after 7 days. CTCF binding was severely impacted at sequence match bound sites (*t-test*, P<10^-5^, 411 sites) along with an increase in H3K9me3 signal (*t-test*, P<10^-5^) (**Supplementary Fig. 7a-c**). This complemented the CTCF loss and H3K9me3 gain results observed with sg4. Therefore, truncated guide CRISPRi can deplete CTCF binding events in a bulk population and at a few hundred genomic loci.

We next explored gene expression effects with CTCF-directed truncated guides. Samples of sg4 and sg8 were collected 7 days post lentiviral transduction and processed for RNA sequencing. Significant genes were determined with dream differential expression analysis ^32^, which aggregates data across replicates for confidence. Gene expression for sg4 indicated no differential genes (**Supplementary Fig. 8**). However, targeting with sg8 resulted in upregulation of 67 genes (**Supplementary Fig. 8**) which may be explained by one or more sg8 sites resulting in genome reorganization due to lost CTCF-mediated loops ^33–36^. Interestingly, the sg8 motif is bound by CTCF at 88.4% of sequence match sites compared to only 46.6% for all tested 10nt guides (**Fig. 3c**). Further investigation is needed to characterize the changes in enhancer-promoter interactions due to CTCF insulator loss. CTCF disruption with one truncated guide led to no differential gene expression while another guide induced gene upregulation.

### Characterization of disrupted CTCF sites

A comprehensive approach to identify significantly impacted CTCF sites was next pursued. We generated a list of over 880k JASPAR CTCF binding sites in the genome (see **Methods**). We plotted CTCF ChIP-seq read counts mapping to these regions for sg4 or sg8 relative to safe harbor controls (**Fig. 4c, Supplementary Figs. 5b** and **7c** and **Supplementary Table 4**). This allowed for proper visualization of both targeted and non-targeted CTCF sites. We then applied dream differential analysis to determine significant sites. This analysis determined that 64% of Jurkat sg4 sites were CTCF depleted (79 out of 123 perfect match bound sites, P<10^-5^, LFC<0.5) (**Fig. 4g, h**). Similarly, dream analysis of sg8 Jurkat samples showed significant loss at 55.7% of CTCF sites (**Supplementary Fig. 7d,** 229 of 411 perfect match bound sites, P<10^-5^, LFC<0.5). We concluded that hundreds of CTCF sites can be depleted with truncated guides and over half of targeted CTCF peaks are lost at this scale.

We next investigated CTCF loss at sites other than perfect match loci. We filtered the 880k JASPAR annotated motifs to find putative CTCF binding sites with ≤9nt complementarity to the 10nt guide that we collectively termed “partial match” sites. We analyzed partial match sites in sg4 containing 1 to 5 mismatches and observed 78 out of 29,908 sites had significant CTCF loss (P<10^-5^, LFC<0.5) (**Fig. 4g, h**). Most partial match sites with CTCF loss had a 1nt mismatch. Motif analysis further illuminated mismatch tolerance positions in the guide at A-T rich bases and the 5’ end for sg4 (**Fig. 4i**) and sg8 (**Supplementary Fig. 7e**). It is noteworthy that most partial match sites remained unaffected by the CRISPRi truncated guides, indicating strong sequence specificity for targeted CTCF sites.

## Discussion

Here we show effective CRISPRi targeting at multiple genomic loci with truncated guides as short as 9nt. This is a unique property of dCas9 moieties, as catalytically active Cas9 is incapable of on-target cleavage with guides shorter than 17nt. A single truncated guide can target CRISPRi to hundreds of TF binding sites, thus expediting the discovery of functional regulatory elements. A library of 24 10nt guides enabled screening of over 13,000 CTCF sites, representing 10.8% and 14% of bound CTCF sites in Jurkat and A375 cells respectively, demonstrating scalable utility. Chromatin binding analysis revealed simultaneous disruption of multiple CTCF binding events with a single truncated guide at most sequence match sites.

The activity of a guide depends on seed sequence, which impacts best outcomes of both 20nt and 10nt guides. When designing a truncated guide library, it is important to consider exact seed sequence as some TF motifs will be better targeted than others. PAM distal mismatches of a few bases are tolerated with full-length guide-directed dCas9 ^37^ and surprisingly we observed the same 5’ flexibility with 10-13nt guides. There are considerations beyond the guide sequence that determine dCas9 targeting efficiency. Not all CREs or TF motifs may be amenable to CRISPRi perturbation. Furthermore, it is unclear how required H3K9me3 is for disruption of CTCF binding. A recent study demonstrated better enhancer and promoter targeting with a KRAB-dCas9-MeCP2 system than with KRAB-dCas9 alone ^38^, suggesting the importance of the repressive marks. In this work we targeted TF binding motifs directly, making it plausible that steric hindrance also contributed to TF displacement from chromatin.

Selection of targeted TF motifs and guide length are critical when planning truncated guide screens. We selected motifs containing an NGG PAM to maximize the likelihood of dCas9 binding. dCas protein variants with expanded PAM requirements will widen the available target sequence space. The shortest amenable truncated guide lengths will need to be determined in these variant systems. In our experiments, 10nt and 13nt guides both significantly disrupted target CTCF binding, however the 13nt guide generated a greater effect size at its targets. Explanations for this are either that longer guides form a more stable R-loop structure with dCas9 resulting in a stronger binding affinity ^39,40^ or longer guides benefit from a more favorable ratio of KRAB-dCas9 protein units to fewer target sites. The maximum number of sites simultaneously targetable with a single guide is yet to be determined. Here we investigate guides with hundreds of sequence match sites (sg4 and sg8), but the exact multi-locus limit will depend on specific guide sequences and expression levels of the KRAB-dCas9 construct. Lastly, we suggest calculating the fraction of sequence match sites bound by the target TF (via ATAC-seq, ChIP-seq, or CUT&Tag data) to assess screen efficiency. In Jurkat cells, 46.6% of sequence match CTCF sites for all guides in the library were bound by the protein. The bound fraction may be lower for other TFs and DNA-binding proteins.

The truncated guide method is a first pass discovery tool for targeting repeated genomic loci. This approach is particularly useful when TF knockout results in negative fitness outcomes or lethality, such as with CTCF. TFs can have alternate cellular functions, such as RNA binding ^41^, that can confound TF knockout studies. This is avoided with truncated guide experiments since binding sites themselves are reliably assayed. Another advantage to this approach is its applicability to cell models with low lentiviral efficiencies or rare cell populations, such as primary cells. A single truncated guide provides a rich landscape of tested outcomes with few transduction events. Finally, we anticipate that truncated guide perturbation will provide a rich readout of gene and TF regulatory networks in single cell assays.

## Methods

### Cell Culture

Jurkat, K562, and MV4-11 cells were cultured in RPMI (Gibco) + 10% FBS (Sigma) supplemented with 1% penicillin/streptomycin (Gibco). HEK293FT and A375 cells were cultured in GlutaMAX High Glucose DMEM (Gibco) + 10% FBS supplemented with 1% penicillin/streptomycin. Cells were tested monthly (negative) for mycoplasma contamination and maintained in a 37°C humidity-controlled incubator with 5% CO_2_. Jurkat, K562, MV4-11 cell lines were obtained from the Cancer Cell Line Encyclopedia (https://portals.broadinstitute.org/ccle/home), HEK293FT cells from Invitrogen, and A375 cells from ATCC. STR profiling was used to confirm cell line identities upon arrival.

### Flow Cytometry

Cells were incubated for 30 minutes at room temperature in 0.5% BSA PBS with 1:50 CD81-FITC antibody (Biolegend, 349504) or mouse IgG1 FITC isotype control antibody (Biolegend, 400107). Cells were washed twice prior to analysis on a Cytoflex (BD) cell analyzer. The gating strategy can be found in **Supplementary Fig. 1e.**

### Guide Selection

All guide sequences in this study can be found in **Supplementary Table 2**. This includes the CTCF motif-directed library for the 24 guides (10nt, selected based on TF motif) and 96 (11nt, by each base to the 5’ end of each 10nt guide). To assess the effect of CTCF knockdown on cell fitness, we used CRISPick to select 20 sgRNAs targeting the CTCF promoter. After screening, one guide was selected based on lethality across all cell lines and was included as the knockdown data found in **Fig 2c**. Additionally, 15 safe harbor sgRNAs ^42^ were included as negative controls.

### Vectors and virus production

Annealed oligos were cloned into an all-in-one KRAB-dCas9-puro vector (pXPR_066, Broad GPP) using the BsmBI restriction enzyme for backbone linearization and T7 ligase for CD81 promoter targeting experiments, the EPB41 enhancer locus experiments, and the CTCF pooled library and follow up. sgCD81i-1 g[9nt] and g[8nt] guides required golden gate assembly (NEB) due to the short length of the oligonucleotides. The CTCF library with varying guide lengths involved the production of 3 separate pooled libraries, one for each of 10nt, 11nt, and 20nt guide lengths. These libraries were then mixed at a balanced (equimolar) ratio to produce the final library.

Single plasmids were chemically transformed into One Shot STBL3 chemically competent coli (Invitrogen C737303). Bacterial cultures were shaken at 225 rpm for one hour at 37°C and then plated on an ampicillin agar dish. After overnight growth at 37°C, single colonies were picked into LB and shaken overnight at 225 rpm and 37°C. Plasmids were isolated the next day using a Plasmid Miniprep Kit (Qiagen) and quantified using a Qubit fluorometer.

Pooled plasmid libraries were electroporated into ElectroMAX™ Stbl4™ electrocompetent cells (Invitrogen 11635018) and spread on to bioassay plates. After overnight incubation at 30°C, bacterial colonies were collected and isolated using the Plasmid Plus Midi Kit (Qiagen 12941). 20nt, 11nt, and 10nt sequences of the CTCF pooled library were cloned as individual pools and then combined to reduce drift. Guide sequences can be found in **Supplementary Table 2**. All guides were cloned with a guanine base at the 5’ end to improve transcription from the U6 promoter ^28,43^.

For single plasmids, 1 x 10^6^ HEK293FT cells were seeded in each 6-well in 2 ml of DMEM + 10% FBS 24 hours prior to transfection. A DNA mixture was prepared consisting of 250µl Opti-MEM, 0.25 µg pCMV_VSVG (Addgene 8454), 1.25 µg psPAX2 (Addgene 12260), 1µg of the all-in-one CRISPR vector (pXPR_066), and 7.5ul TransIT-LT1 (Mirus) transfection reagent. After a 20-minute incubation, the solution was added dropwise to the 6-well and incubated for 6 – 8 hours. Fresh media was added to the cells and collected 36 hours later and either snap frozen or added to cells.

For CTCF pooled library, 8 x 10^6^ HEK293FT cells were seeded in each of 2 T75 flasks in 12 ml of DMEM + 10% FBS 24 hours prior to transfection. Next, pCMV_VSVG (Addgene, 8454, 1.5 µg), psPAX2 (Addgene 12260, 9 µg), the guide containing vector (pXPR_066, 7.5 µg), and 66 µl TransIT-LT1 (Mirus MIR 2306) were combined with 2.1 ml of Opti-MEM to produce TransIT-LT1:DNA complexes. After a 20-minute incubation, the solution was added dropwise to the 6-well and incubated for 6 – 8 hours, then the media was changed. After 36 hours, the lentivirus was collected, filtered, and either snap frozen or used for cell transduction.

### Viral transduction

CTCF pooled library frozen viral supernatant (300 µl) was thawed and added to 700 µl target cells in 12-wells with a final volume of 10 µg/mL polybrene, resulting in a 30–50% transduction efficiency, corresponding to an MOI of ∼0.35–0.70. Cells with viral supernatant were centrifuged at 2000 x g for 20 minutes at 22°C and incubated overnight. After 18-24 hours, cells were fed fresh media and maintained at 2 x 10^5^ cells/mL for suspension cells (Jurkat, K562, and MV4-11) and 1-2 x 10^5^ cells/cm^2^ for adherent cells (A375 and HEK293). Cells were passaged into media supplemented with 1 µg/mL puromycin 3 days after transduction. Seven days after transduction, cells were passaged into 0.5 µg/mL puromycin (1/2 dose) and cultured continuously for the duration of the screen. At day 21, pellets of 1 x10^6^ cells were snap frozen on dry ice and stored at −80°C in preparation for gDNA isolation.

Single transductions were performed identically to the pooled production, with the exception that viral supernatant varied based on viral titer.

### Genomic DNA preparation and sequencing

Genomic DNA was isolated using DNeasy Blood and Tissue Kit (Qiagen 69504). PCR, sequence adaptor barcoding, cleanup, sequencing, and data deconvolution were carried out as previously described ^44^. PCR primers were Argon and Beaker (Broad Institute GPP). At the PCR stage, CTCF pooled library plasmid DNA (pDNA) was diluted to 10ng for amplification. All PCR reactions were carried out for 28 cycles. Libraries were prepared using TruSeq amplicon construction and single end sequenced on a MiSeq50. Fastq files were deconvolved using PoolQ (https://portals.broadinstitute.org/gpp/public/software/poolq). Apron (Broad Institute GPP) was used to analyze the distribution of each guide relative to the plasmid DNA, enabling enrichment/depletion measurements.

### ChIP-seq sample preparation

Frozen crosslinked cell pellets (1 x 10^7^ cells) were suspended in cell lysis buffer (20 mM Tris pH 8.0, 85 mM KCl, 0.5% NP40) with protease inhibitors (cOmplete EDTA-free Protease Inhibitor Tablets, Sigma Aldrich), incubated on ice for 10 min, then centrifuged at 1000 x g for 5 minutes. Cell pellets were resuspended for a second time in cell lysis buffer with protease inhibitors, incubated on ice for 5 minutes and centrifuged for 5 minutes at 1000 x g. The pellets were resuspended in nuclear lysis buffer (10 mM Tris-HCl pH7.5, 1% NP40, 0.5% sodium deoxycholate, 0.1% SDS) with protease inhibitors for 10 min and subsequently sheared in a sonifier (Branson).

The chromatin was quantified after sonication to determine the cell number in each sample. H3K9me3 ChIP-seq samples were prepared with 1.5 x 10^6^ cells and 0.4 µg H3K9me3 antibody (Abcam ab176916). CTCF ChIP-seq samples were prepared with 3 x 10^6^ cells and 1 µg CTCF antibody (Diagenode C15410210). ChIP-seq Dilution Buffer (16.7 mM Tris-HCl pH 8.1, 167 mM NaCl, 0.01% SDS, 1.1% Triton X-100, 1.2 mM EDTA) with protease inhibitors was added to bring the ChIP volume to 0.5 mL. ChIP-ses samples were rotated overnight at 4°C. The following day, Protein A Dynabeads (Invitrogen) were added for 1 hour to enrich fragments of interest. The ChIP-seq samples were removed from rotation and placed on a magnet to isolate the beads. The beads were washed with a series of buffers, low salt RIPA buffer, high salt RIPA buffer, LiCl buffer (250mM LiCl, 0.5% NP40, 0.5% sodium deoxycholate, 1mM EDTA,10mM Tris-HCl pH 8.1) and finally Low TE. The Protein A beads were then suspended in 50 µl elution buffer (10 mM Tris-Cl pH 8.0, 5 mM EDTA, 300 mM NaCl, 0.1% SDS and 5 mM DTT directly before use) and 8 µl of reverse crosslinking mix (250mM Tris-HCl pH 6.5, 1.25 M NaCl, 62.5 mM EDTA, 5 mg/ml Proteinase K, and 62.5 µg/ml RNAse A). The suspended beads were incubated at 65°C for a minimum of 3 hours. After incubation, the supernatants were transferred to a clean tube. The DNA was SPRI purified, eluted, and quantified by Qubit. Libraries with 6 µg of input were prepared using the KAPA Hyper Prep Kit.

### Quantitative real time PCR

Real time PCR was performed as described previously ^45^. In brief, RNA extraction was performed with the RNeasy Plus Micro Kit (Qiagen) and cDNA was synthesized using the Superscript III First-Strand Synthesis System for RT-PCR (Invitrogen). Probes for EPB41 and actin beta transcripts (EPB41_1_For AACTTCCCAGTTACCGAGCA, EPB41_1_Rev CTTGAGTCCGGCCACTGTAT, EPB41_2_For CTGCTCTAGTGGCCTTCTGG, EPB41_2_Rev CTGCTCGGTAACTGGGAAGT, actin-b_For CATCGAGCACGGCATCGTCA, and actin-b_Rev TAGCACAGCCTGGATAGCAAC) were paired with Power SYBR Green PCR Master Mix (Applied Biosystems) for quantification. Samples were analyzed on a BioRad CFX Opus 384 Real-Time PCR System. All samples were normalized to the average Ct across all replicates of actin beta safe harbor.

### RNA sequencing sample preparation and analysis

Jurkat and A375 cells were transduced with all-in-one KRAB-dCas9-puro (pXPR_066) vector carrying safe harbor, CD81 CRISPRi (20nt or g[9nt]), CTCF-sg4 (10nt), or CTCF-sg8 (10nt) guides (**Supplementary Table 2**) were pelleted and stored in −80°C. RNA was isolated with RNeasy Plus Micro kit (Qiagen) according to the manufacturer’s protocol and ensuring RIN values greater than 7. Libraries were prepared first with Poly-A enrichment using magnetic oligo(dT)-beads (Invitrogen), then ligated to RNA adaptors for sequencing. Paired end sequencing (2 x 150bp) was carried out on an Illumina Nextseq or Novaseq (Illumina).

### Data analysis

#### Pooled screening

Log fold change calculations were calculated with the starting plasmid DNA pool as reference. Initial quality control of pooled screening data included running pairwise comparisons on replicates to assess replicate consistency. Based on these tests, one replicate of the screen in K562 cells was excluded, as multiple comparisons test of these replicates identified a significant difference between replicate LFC values (Repeated measures one-way ANOVA, P<0.0001) and a post-hoc multiple comparisons test identified significant differences between Rep A and Rep B as well as between Rep B and Rep C (Tukey’s, P<0.0001 for both tests) while there was no significant difference between Rep A and Rep C (Tukey’s, p=0.794). Based on these findings, Rep B was excluded from further analysis while the 2 other replicates from this screen were retained. All other cell line replicates were not significantly different.

Z-scores for pooled screen analysis were calculated using the following equation:\

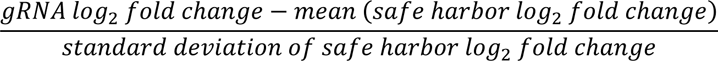

#### RNA-seq data processing

RNA-seq data for A375 sgCD81, Jurkat sg4 and sg8 were processed using the Kallisto v0.46.1 alignment and quantification tool (*kallisto quant -i {transcriptome_index} -o output -b 50 ∼{read1}∼{read2} -t 4 -g {gtf}*). Transcriptome index and gtf for human was taken from https://github.com/pachterlab/kallisto-transcriptome-indices/releases. The data, composed of paired-end reads in fastq format and included multiple replicates per condition. Output files of interest to this analysis were the .h5 count matrices and alignment logs.

#### RNA-seq quantification and analysis

The .h5 count matrices were imported into R using *tximport::tximport*. Raw counts were aggregated to the gene level. Genes with zero counts across all samples were subsequently removed from the matrix for differential expression analysis. Differential expression followed the workflow outlined in the vignette provided within the *dream* ^32^ statistical package. To visualize the impact of the guide in comparison to the Safe Harbor, we generated volcano plots (**Fig. 1e** and **Supplementary Fig. 8a, b**). We set the significance criteria to have an absolute log fold-change (abs(logFC)) > 2 and an adjusted P<0.001. The top differentially expressed genes were visualized through heatmaps using *pheatmap::pheatmap*, with hierarchical clustering revealing that replicates clustered together.

#### Comparison of CTCF guide target sites and CTCF binding events

Perfect match sites were identified using Cas-OFFinder v2.4 ^46^ (http://www.rgenome.net/cas-offinder/) with hg38 2bit as reference. Alt chromosomal matches were excluded from the analysis. “N” bases were added to guide sequences such that they met the minimum threshold of 15 nt.

#### Putative CTCF binding site determination

CTCF sites were selected using the JASPAR MA0139 matrix profile (https://jaspar.genereg.net/matrix/MA0139.1/) and filtered down to the 880k sites using the R library and steps previously detailed ^47^. These were exported to a .bed file and a .saf file.

#### ChIP-seq data processing

For **Fig. 3c** and **Supplementary Fig. 3a, b,** Jurkat and A375 CTCF ChIP-seq datasets were processed using the ENCODE ChIP-seq pipeline v2.1.5 (https://github.com/ENCODE-DCC/chip-seq-pipeline2). Both replicates were processed using the default “tf” options for the pipeline with the MACS2 peak-caller. To obtain a final peak-set of CTCF binding events, we utilized the IDR-optimal output peak calls at an IDR threshold of < 0.05 for each cell line and merged overlapping peaks. We then extended the CTCF peaks symmetrically by +/- 50 bp, corresponding to a stringent perturbation radius, and used bedtools to obtain overlapping sites between each CTCF guide target site and CTCF binding peaks.

For all other figures, ChIP-seq data for CTCF and H3K9me3 were processed using the ENCODE ChIP-seq pipeline v2.2.0 with default parameters. The *pipeline_type* parameter for CTCF and H3K9me3 was set to “tf” and “histone”, respectively. The data, composed of single-end reads in fastq format, included two replicates per condition. Output files of interest to this analysis were the bam files and QC html reports.

#### ChIP-seq quantification and analysis

CTCF bigwig files were created using bamCoverage (*bamCoverage -b $1 -o “$2.bw” -bs 50 -p 4 --effectiveGenomeSize 2913022398 --normalizeUsing bpm*). We used deeptools to calculate normalized signal using counts within a +/- 3Kb window centered at perfect match sites. To observe the effect of the guide in the H3K9me3 landscape, we extracted the windows overlapping perfect match sites and plotted the aggregate histone signal in Safe Harbor and sg4 samples **(Fig. 4d)**. We also generated CTCF profile heatmaps (**Supplementary Fig. 5a).** The bigwigs were used with the *karyoploteR* R package to generate genome tracks **(Fig. 4e, f)**

For visualization and differential binding analysis (**Fig. 4** and **Supplementary Figs. 5b, 6, and 7**), we created a CTCF count matrix using featureCounts (*featureCounts(files, allowMultiOverlap = T, largestOverlap = T, annot.ext = “jaspar_motifs.saf”, readExtension3 = 200, ignoreDup = T)*, where *jaspar_motifs.saf* contains the putative CTCF binding sites with window size 500bp in a .saf format. The same was done for a H3K9me3 count matrix, except the .saf file contains genome-wide non-overlapping windows of size 5Kb. Raw counts were stored in a .tsv file. To observe the effect of the guide in CTCF binding at perfect match sites, we first loaded the count matrix in R (rows: ∼880k putative binding sites from JASPAR, columns: samples) and used *edgeR::cpm* to normalize the data. We extracted the bins that overlapped a perfect match site and filtered out bins if they had <5 CPM in the Safe Harbor samples. CTCF binding was averaged between sample replicates. We observed a significant difference in mean CTCF binding between Safe Harbor and sg4 samples, using *stats::t.test* (**Fig. 4a, Supplementary Figs. 6, 7a, 7b)**. A similar procedure was used for H3K9me3 (**Fig. 4b)**.

To observe the genome-wide effect of the guide in CTCF binding, we took the above normalized CTCF count matrix and filtered out bins if they had <5 CPM in the Safe Harbor samples. We plotted the counts in the remaining bins and colored points if they overlap a sg4 or aag[sg4] perfect match site (**Fig. 4c, Supplementary Figs. 5b, 6, 7d).**

For the differential analysis, we utilized the *dream* statistical package in R, as outlined by Hoffman and Roussos, 2021. We established significance criteria with a requirement for absolute log fold change (abs(logFC)) > 0.5 and an adjusted P<10^-5^. (**Fig. 4g, Supplementary Fig. 7e).** While many significant sites overlapped perfect match sites, we also observed some sites that showed significant CTCF loss where the sequence did not perfectly match the guide target sequence. We extracted the sequence at these sites from JASPAR MA0139, and used *Biostrings::consensusMatrix* and *ggseqlogo:: ggseqlogo* to look at the logogram of the sequences **(Fig. 4i, Supplementary Fig 7e)**. We observed that the sequences matched closely with the guide target sequence. Further analysis showed most of these sequences had only 1 or 2 mismatches from the target sequence **(Fig 4h)**.

R (version 4.1.2), Python (version 3.7), and Graphpad Prism (version 10) were used for visualization.

## Data Availability

Raw ChIP-seq and RNA-seq data will be available online after publication.

## Code Availability

Link to code/scripts here: https://github.com/broadinstitute/gro-crispri-ctcf (available after publication)

## Acknowledgements

We thank E. Gaskell, N. Durand, A. Hall, A. Cruz, and C. White for helpful discussions, and E. Donnard and E. Roberts for reagents. The Broad Institute Flow Core and Genomics Platform provided experimental support. This project was supported by funds from the Gene Regulation Observatory at the Broad Institute. B.E.B. is the Richard and Nancy Lubin Family Endowed Chair at the Dana Farber Cancer Institute and an American Cancer Society Research Professor.

## Author Contributions

M.M.M, R.I., A.C., and Y.L. conducted the experiments. S.W., N.J., and E.M. processed the data. J.D.B, C.B.E, J.G.D., B.E.B., N.S. and F.J.N. provided guidance and direction. M.M.M. and F.J.N. conceived the study and wrote the paper with help from all co-authors.

## Supplementary Table Legends

**Supplementary Table 1.** RNAseq counts matrices. Gene level normalized counts for samples. A375 samples included untreated or treated with Safe Harbor or CD81 guides Jurkat cells included untreated or treated with Safe Harbor, sg4 or sg8.

**Supplementary Table 2.** Guides table. A listing of all CRISPR guide sequences and respective oligonucleotides used in this study.

**Supplementary Table 3.** CTCF screening data. Jurkat, A375, MV4-11, HEK293 and K562 cells transduced with CTCF library and processed after 21 days. Data are guide representation depicted by log_2_-fold change as normalized to plasmid DNA and z-score as normalized to safe harbor guides.

**Supplementary Table 4.** CTCF site quantification. Count matrix of 880k potential CTCF binding sites (rows) and samples (columns). Data are raw counts from CTCF ChIP and 500bp bin size of the putative binding sites.

