## Supplementary_Figures for "Multi-locus CRISPRi targeting with a single truncated guide RNA"

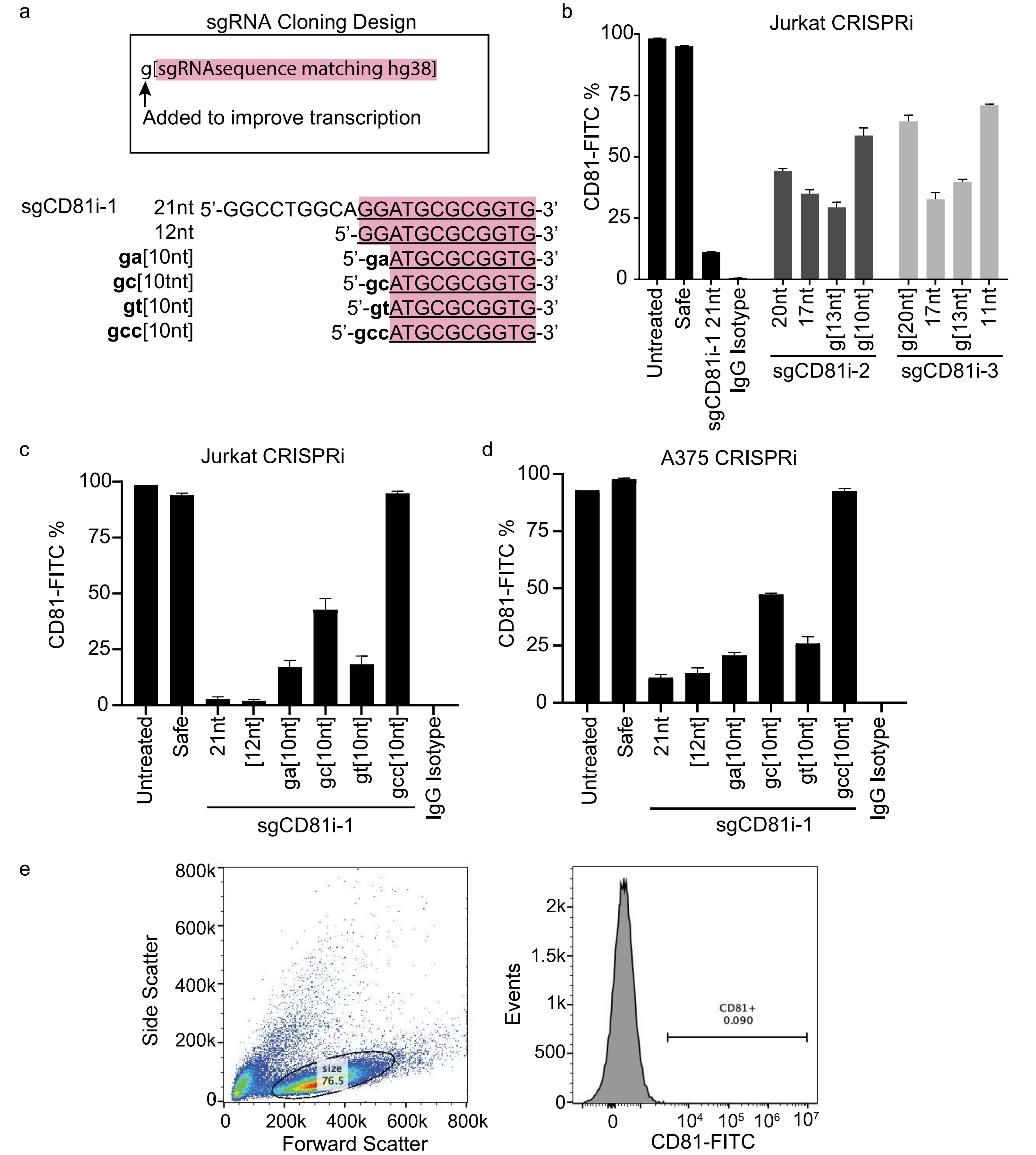


**Supplementary Figure 1.** CRISPRi performance with additional truncated guides. **a** Convention for guide cloning design with target matching sequence highlighted in pink. Guide sequences tested are based on 5' truncations in sgCD81i-1, that targets the CD81 TSS. **b** CRISPRi treatment of two additional CD81 TSS targeting guides and respective 5' truncations. CD81 expression was read by flow cytometry. **c**, **d** Jurkat (**c**) and A375 (**d**) cells CRISPRi treated with the CD81i-1 based guides in panel **a** and analyzed for CD81 expression by flow cytometry. All data in **b-d** are triplicate with mean +/- SEM plotted. **e** Gating strategy for live cell selection (left panel) and CD81+ selection by setting the gate <0.1% positive in untreated isotype control stained cells (right panel).

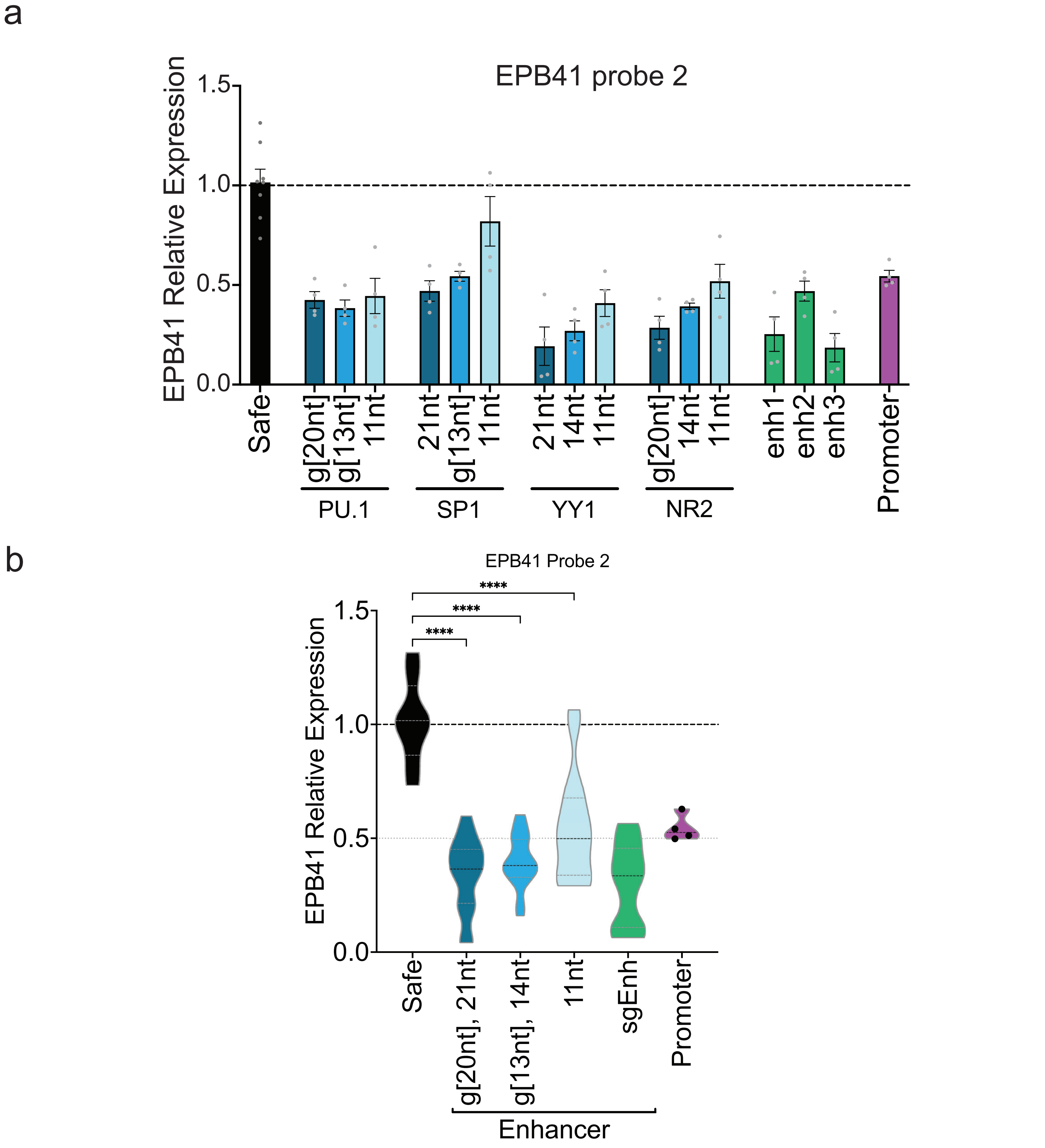


**Supplementary Figure 2.** Alternate qPCR probe for EPB41 expression. **a** Quantitative real time-PCR analysis of EPB41 expression with respective CRISPRi guide treatment conditions in K562 cells. Promoter, enhancer guides (validation set: enh1, enh2, enh3, Gasperini et al. 2019), and Safe (Safe Harbor targeting) are all 20nt guides. Data are normalized to mean value of Safe Harbor actin-b control probe and mean +/- SEM. **b** Violin plot summary of CRISPRi targeting data by guide length or targeting region category. Results are analyzed by one-way ANOVA. p****<0.0001 for all guide lengths compared with Safe Harbor guide.

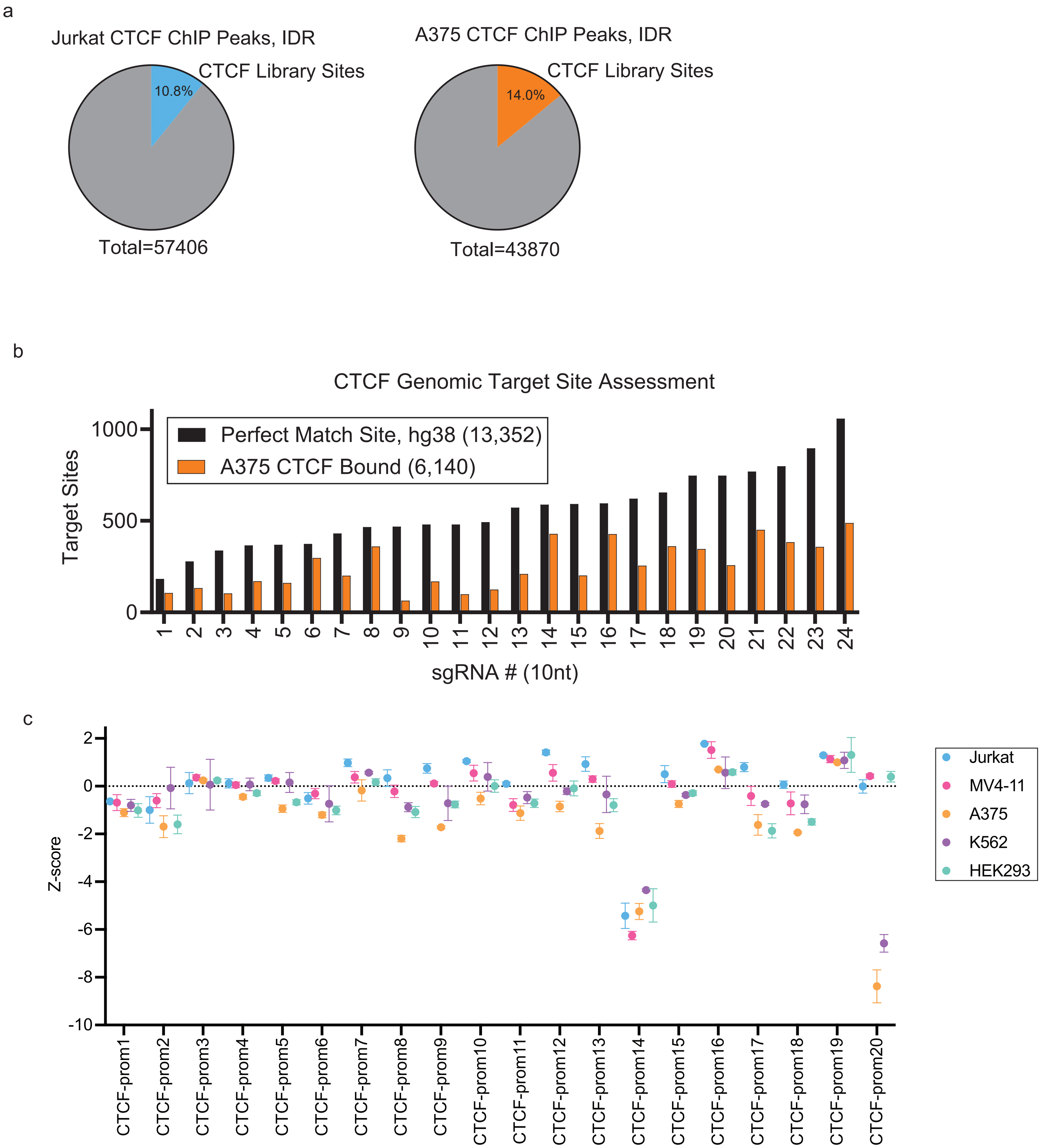


**Supplementary Figure 3.** CTCF library analysis. **a** Fraction of CTCF ChIPseq sites targeted by the 24 guide library in Jurkat (left, blue) and A375 (right, orange) cells. **b** CTCF ChIPseq data depicting the fraction of perfect match sites with CTCF bound in A375 (orange) cells for each guide in the CTCF library. **c** CTCF promoter targeting guide fitness in the 5 cell lines tested (indicated). Pooled screens were run in triplicate, while K562 was in duplicate. Data are mean +/- SD.

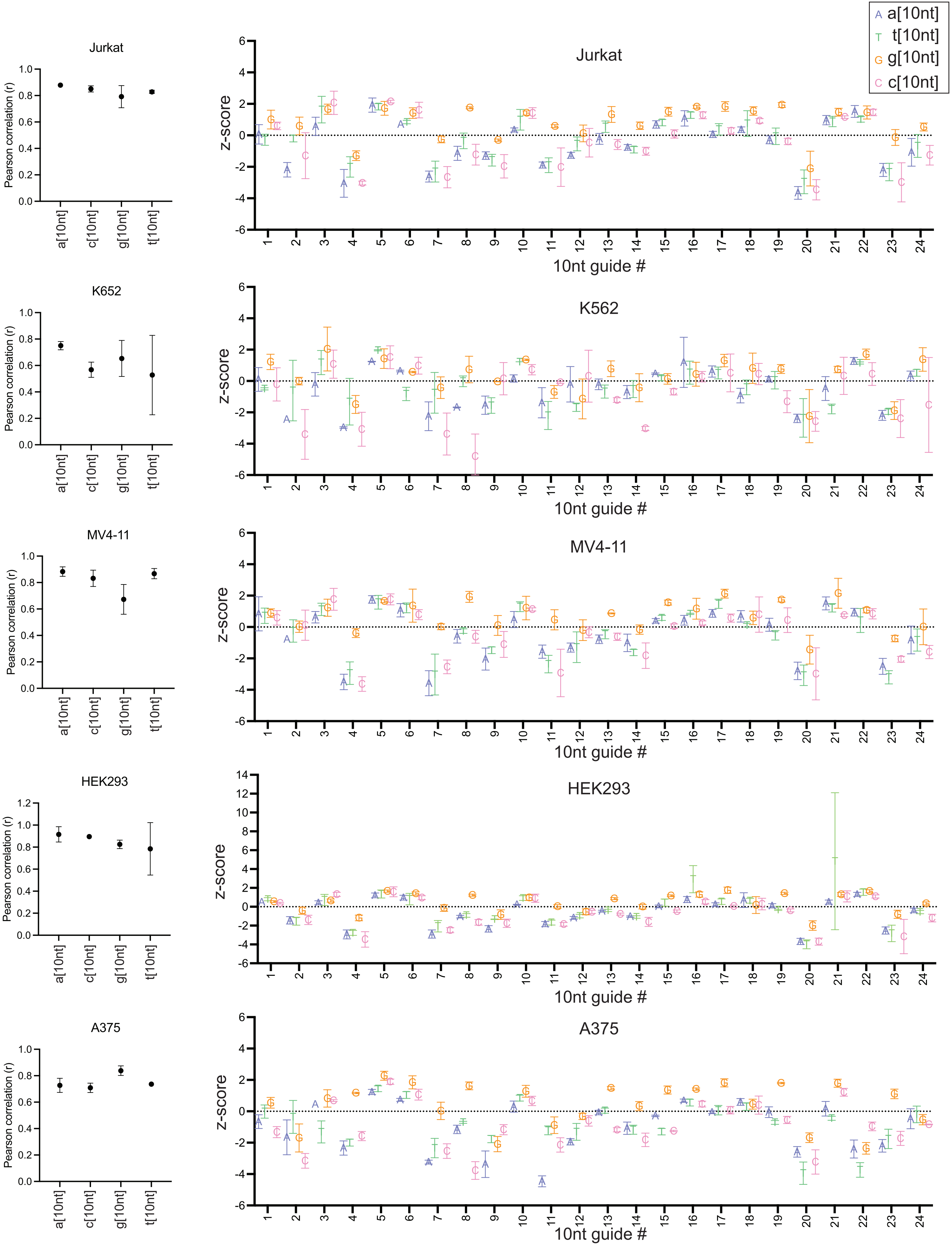


**Supplementary Figure 4.** Appending an 11th nucleotide to the CTCF library guides. Left, pearson correlations for the n[10nt] screening data compared to the respective 10nt guide data for the indicated cell line. Right, fitness effects of each n[10nt] guide for each of the cell lines tested. z-score calculated from log fold change relative to safe harbor control. Experiments were run in triplicate except for K562, in duplicate.

a

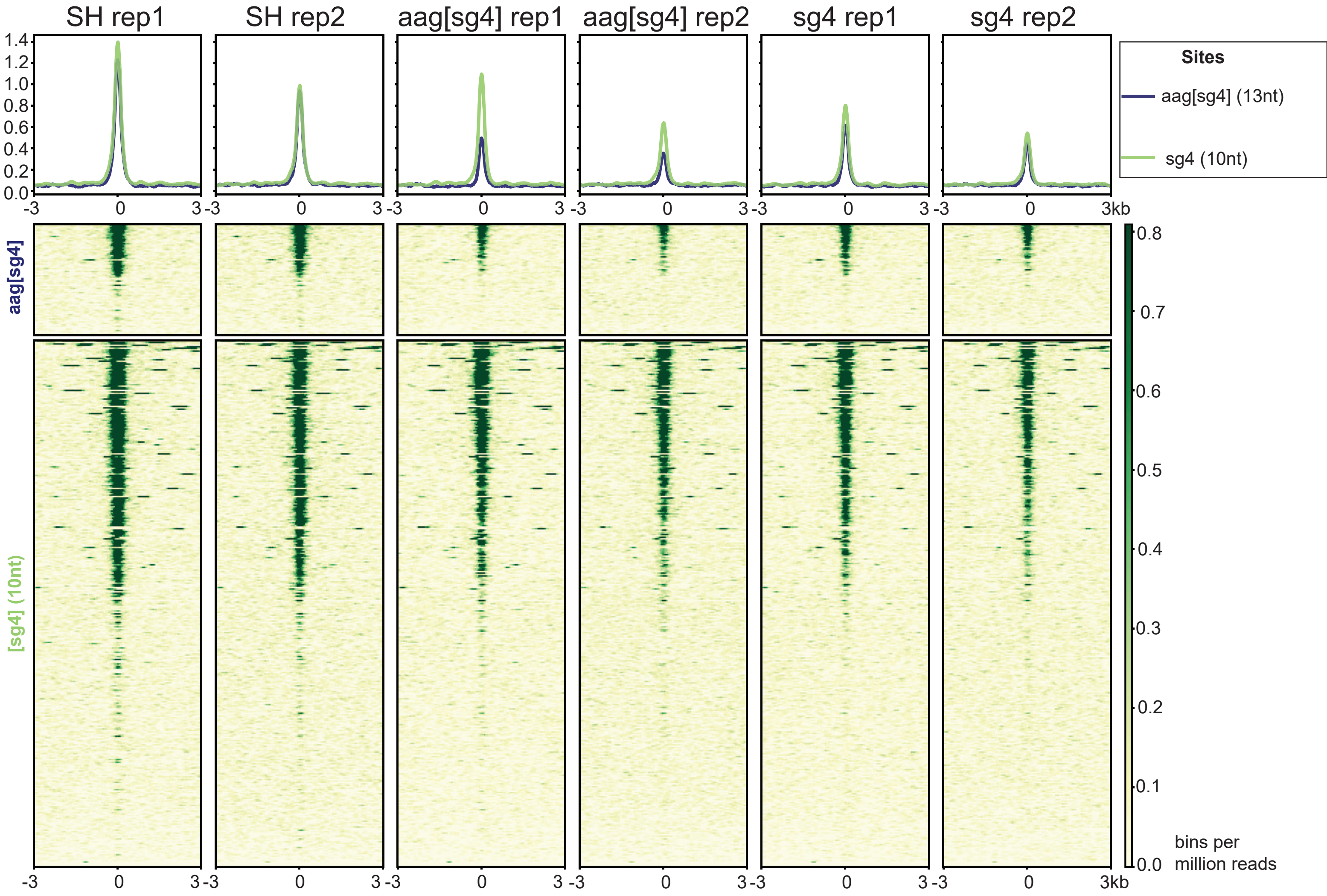


b

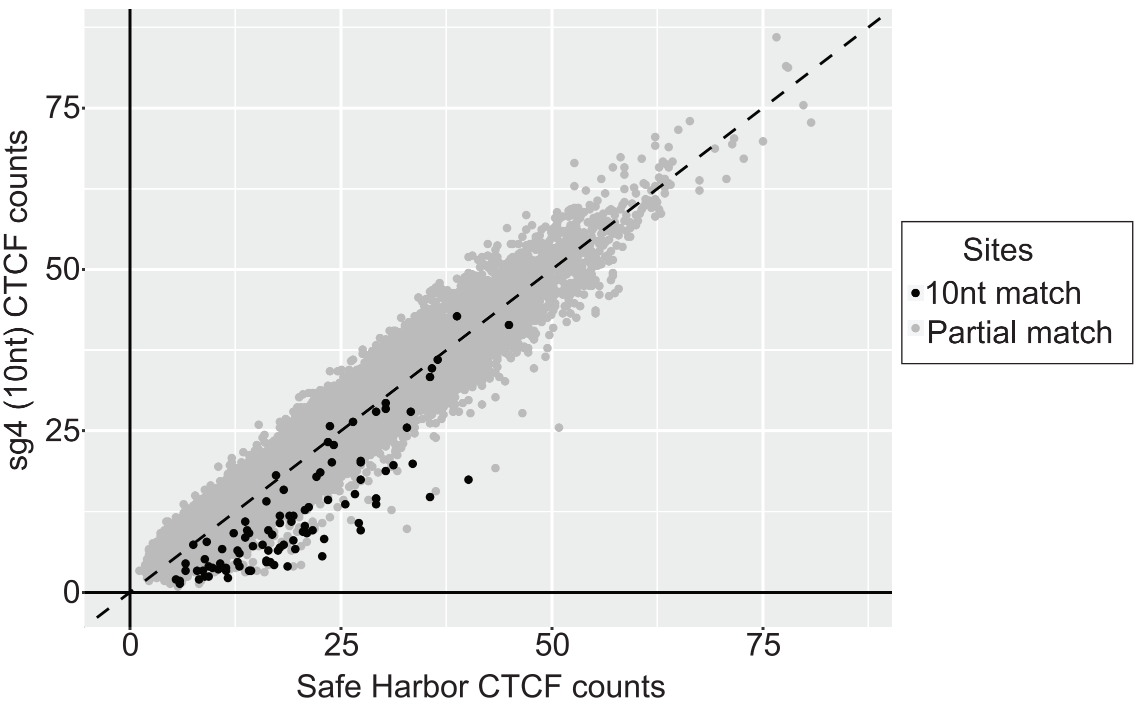


**Supplementary Figure 5.** Specificity and selectivity of sg4 and aag[sg4] guides. **a** CTCF ChIPseq density plot and heatmaps of sg4 and aag[sg4] targeted loci with a 6kb window for Jurkat cells treated with sg4, aag[sg4], or Safe Harbor (SH) guides. Top heatmap cluster represents sites with an aag[sg4] sequence match, which sg4 also matches, and the bottom heatmap cluster are sites only sg4 has a sequence match. **b** Scatter plot of 880k JASPAR annotated CTCF sites, rep2. Rep1 is found in **Fig. 4c**.

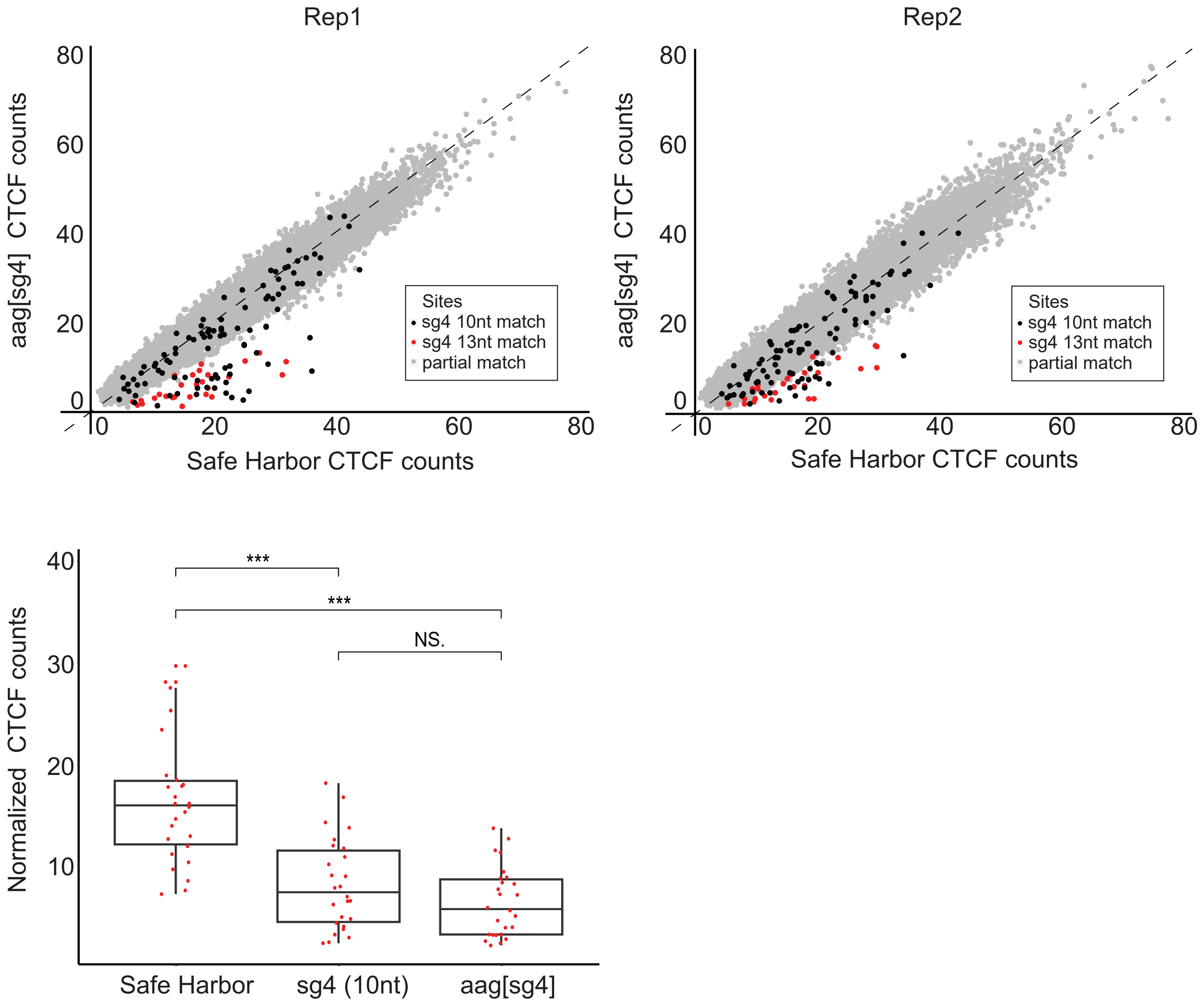


**Supplementary Figure 6.** CTCF targeting with aag[sg4]. **Top,** scatter plots of CTCF binding at JASPAR motifs (880k) with perfect match sites (black), aag[sg4] (13nt) sites (red) partial match sites (grey). **Bottom**, boxplot of CTCF binding at 26 genomic loci targeted by both sg4 and aag[sg4]. t-test ***P=2.16 x $10^{-6}$ for sg4 and ***P=1.48 x $10^{-8}$ for aag[sg4]. Comparison between sg4 and aag[sg4] was not significant (NS).

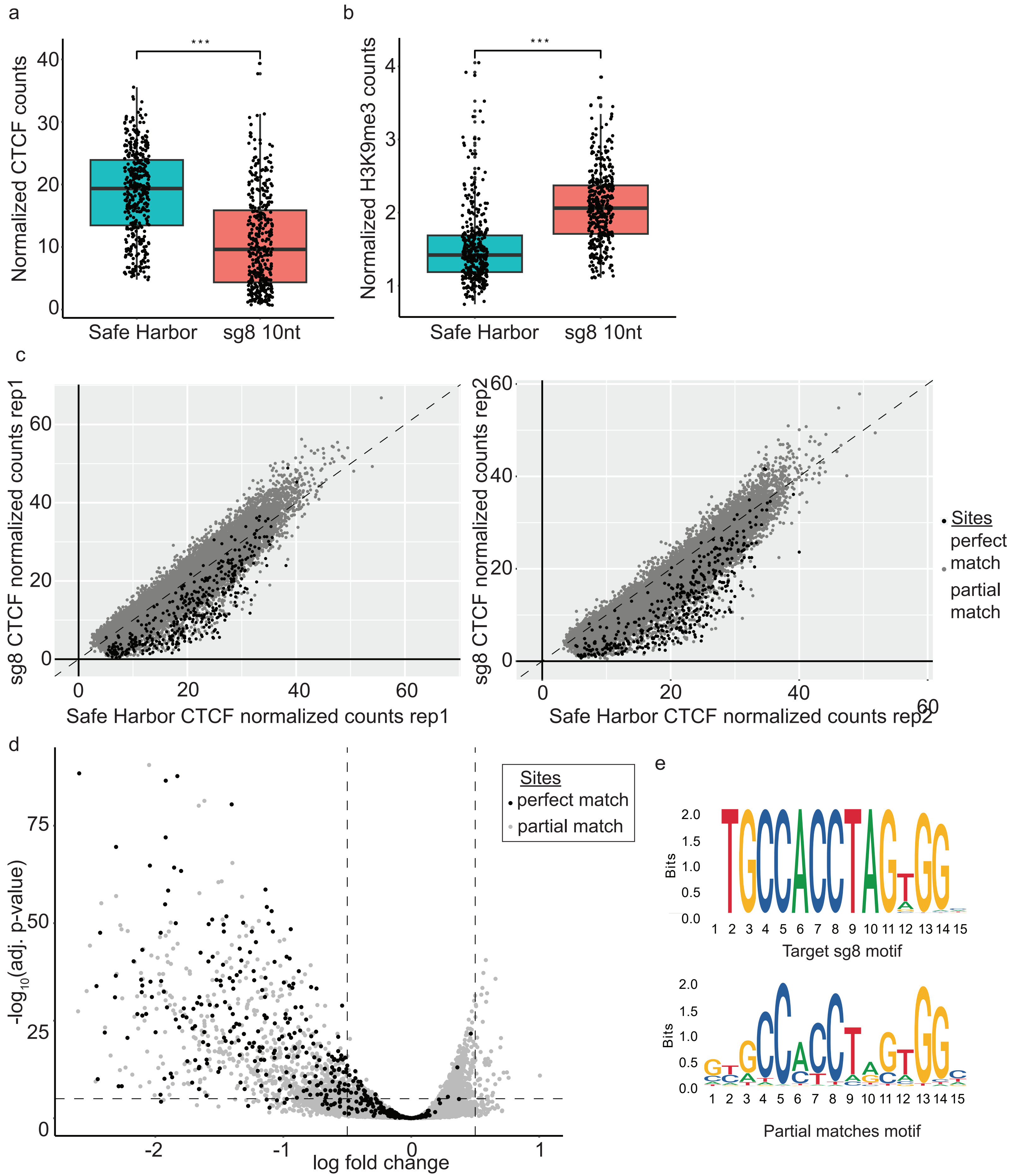


**Supplementary Figure 7.** sg8 targeting data. **a** Boxplot of CTCF signal at bound sequence match sites of sg8 and safe harbor treated Jurkat cells. Significance with t-test ***P=5.02 x $10^{-47}$. **b** Boxplot of H3k9me3 signal at sequence match sites. Significance with t-test ***P=4.43 x $10^{-46}$. **c** Scatter plot of JASAR-filtered CTCF sites with sg8 perfect match sites (black) and all other sites (grey) in rep1 and rep 2 Jurkat cells. **d** Significantly targeted sites determined by Dream analysis with the same annotations as in **c**. **e** Perfect match motif (top) and partial match sites motif (bottom) with sg8.

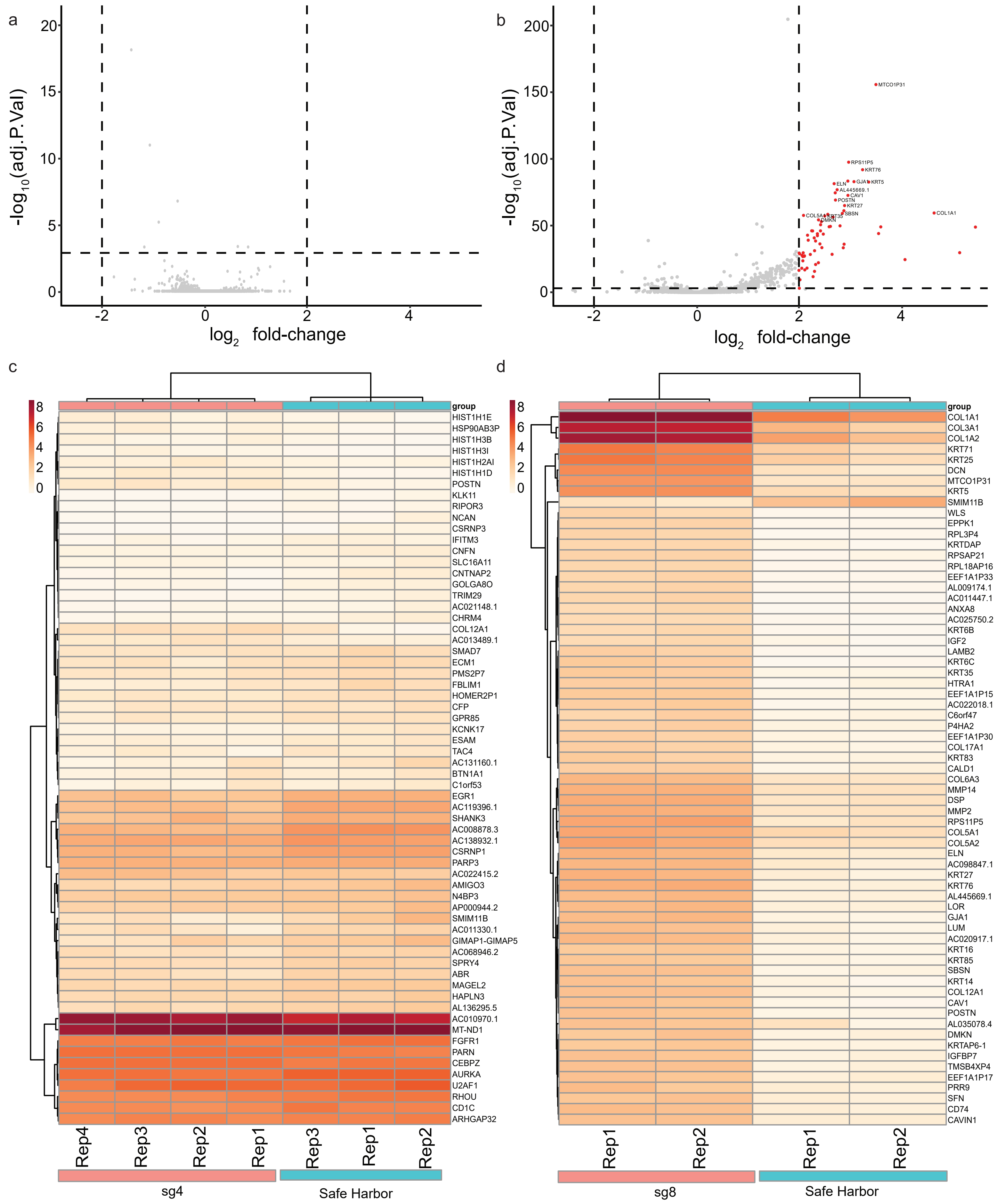


**Supplementary Figure 8.** Gene expression in CTCF truncated guides. Jurkat cells treated with sg4, sg8, or safe harbor targeting CRISPRi lentivirus for 7 days and processed by RNAseq. **a, b** Volcano plots of sg4 (**a**) and sg8 (**b**) normalized to safe harbor. Dotted lines represent thresholds set at $\log_2$ fold change [abs 2] and -log(P-value)>3. **c, d** Heatmaps of the top differential genes in sg4, none signficant (**c**) and the top 67 significant differential genes in sg8 (**d**) alongside safe harbor control gene expression. See **Methods** for details.
